# *Severe acute respiratory syndrome-related coronavirus*: The species and its viruses – a statement of the Coronavirus Study Group

**DOI:** 10.1101/2020.02.07.937862

**Authors:** Alexander E. Gorbalenya, Susan C. Baker, Ralph S. Baric, Raoul J. de Groot, Christian Drosten, Anastasia A. Gulyaeva, Bart L. Haagmans, Chris Lauber, Andrey M Leontovich, Benjamin W. Neuman, Dmitry Penzar, Stanley Perlman, Leo L.M. Poon, Dmitry Samborskiy, Igor A. Sidorov, Isabel Sola, John Ziebuhr

**Affiliations:** Departments of Biomedical Data Sciences and Medical Microbiology, Leiden University Medical Center, Leiden, The Netherlands; Faculty of Bioengineering and Bioinformatics and Belozersky Institute of Physico-Chemical Biology, Lomonosov Moscow State University, 119899 Moscow, Russia; Department of Microbiology and Immunology, Loyola University of Chicago, Stritch School of Medicine, Maywood, Illinois, USA; Department of Epidemiology, University of North Carolina, Chapel Hill, North Carolina, USA; Division of Virology, Department of Biomolecular Health Sciences, Faculty of Veterinary Medicine, Utrecht University, Utrecht, The Netherlands; Institute of Virology, Charité - Universitätsmedizin Berlin, Berlin, Germany; Viroscience Lab, Erasmus MC, Rotterdam, The Netherlands; Texas A&M University-Texarkana, Texarkana, TX, USA; Department of Microbiology and Immunology, University of Iowa, Iowa City, Iowa, USA; Centre of Influenza Research & School of Public Health, The University of Hong Kong, Hong Kong, People’s Republic of China; Department of Molecular and Cell Biology, National Center of Biotechnology (CNB-CSIC), Campus de Cantoblanco, Madrid, Spain; Institute of Medical Virology, Justus Liebig University Giessen, Giessen, Germany

**Keywords:** Coronaviruses, comparative genomics, virus evolution, nomenclature, phylogenomics, respiratory distress syndrome, species, taxonomy, virus, zoonosis

## Abstract

The present outbreak of lower respiratory tract infections, including respiratory distress syndrome, is the third spillover, in only two decades, of an animal coronavirus to humans resulting in a major epidemic. Here, the Coronavirus Study Group (CSG) of the International Committee on Taxonomy of Viruses, which is responsible for developing the official classification of viruses and taxa naming (taxonomy) of the *Coronaviridae* family, assessed the novelty of the human pathogen tentatively named 2019-nCoV. Based on phylogeny, taxonomy and established practice, the CSG formally recognizes this virus as a sister to severe acute respiratory syndrome coronaviruses (SARS-CoVs) of the species *Severe acute respiratory syndrome-related coronavirus* and designates it as severe acute respiratory syndrome coronavirus 2 (SARS-CoV-2). To facilitate communication, the CSG further proposes to use the following naming convention for individual isolates: SARS-CoV-2/Isolate/Host/Date/Location. The spectrum of clinical manifestations associated with SARS-CoV-2 infections in humans remains to be determined. The independent zoonotic transmission of SARS-CoV and SARS-CoV-2 highlights the need for studying the entire (virus) species to complement research focused on individual pathogenic viruses of immediate significance. This research will improve our understanding of virus-host interactions in an ever-changing environment and enhance our preparedness for future outbreaks.

## Is the human coronavirus that emerged in Asia in December 2019 novel?

Is the outbreak of an infectious disease caused by a new or a previously known virus (**Box 1**)? This is among the first and principal questions because the answer informs measures to detect the causative agent, control its transmission and limit potential consequences of the epidemic. It also has implications for the virus name. On a different time scale, the answer also helps to define research priorities in virology and public health.

The questions of virus novelty and naming are now posed in relation to a coronavirus causing an outbreak of a respiratory syndrome that was first detected in Wuhan, China, December 2019. It was temporally named 2019 novel coronavirus, 2019-nCoV. The term “novel” may refer to the disease (or spectrum of clinical manifestations) that is caused in humans infected by this particular virus, which, however, is only emerging and requires further studies^1,2^. The term “novel” in the name of 2019-nCoV may also refer to an incomplete match between the genomes of this and other (previously known) coronaviruses, if the latter was considered an appropriate criterion for defining “novelty”. However, virologists agree that neither the disease nor the host range can be used to reliably ascertain virus novelty (or identity), since few genome changes may attenuate a deadly virus or cause a host switch^3^. Likewise, we know that RNA viruses persist as a swarm of co-evolving closely related entities (variants of a defined sequence, haplotypes), known as quasispecies^4,5^. Their genome sequence is a consensus snapshot of a constantly evolving cooperative population *in vivo* and may vary within a single infected person^6^ and over time in an outbreak^7^. If the strict match criterion of novelty was to be applied to RNA viruses, it would have qualified every virus with a sequenced genome as a novel virus, which makes this criterion poorly informative. To get around the potential problem, virologists instead may regard two viruses with non-identical but similar genome sequences as variants of the same virus; this immediately poses the question of how much difference is large enough to recognize the candidate virus as novel or distinct? This question is answered in best practice by evaluating the degree of relatedness of the candidate virus to previously known viruses of the same host or established monophyletic groups of viruses, often known as genotypes or clades, which may or may not include viruses of different hosts. This is formally addressed in the framework of virus taxonomy (**Box 2**).

In this study, we present an assessment of the novelty of 2019-nCoV and detail the basis for (re)naming this virus severe acute respiratory syndrome coronavirus 2, SARS-CoV-2, which will be used hereafter.

## Defining novelty and the place of SARS-CoV-2 within the taxonomy of the *Coronaviridae* family

During the 21^st^ century, researchers studying coronaviruses – a family of enveloped positive-stranded RNA viruses of vertebrates^8^ – were confronted several times with the question of coronavirus novelty, including two times when a severe or even life-threatening disease was introduced into humans from a zoonotic reservoir: this happened with severe acute respiratory syndrome (SARS)^9-12^ and, a few years later, with Middle East respiratory syndrome (MERS)^13,14^. Each time, the pathogen was initially called a new human coronavirus, as was the case with SARS-CoV-2 during the current outbreak, every time the issue was resolved by the sequence-based family classification.

The current classification of coronaviruses includes taxa at eight out of the fifteen available ranks^15^, and it recognizes forty-nine species in twenty-seven subgenera, five genera and two subfamilies that belong to the family *Coronaviridae*, suborder *Cornidovirineae*, order *Nidovirales*, realm *Riboviria*^16-18^. The family classification and taxa naming (taxonomy) are developed by the Coronavirus Study Group (CSG), a working group of the International Committee on Taxonomy of Viruses (ICTV)^19^. The CSG has responsibility in assessing the novelty of viruses through their relation to known viruses in established taxa and, for the purpose of this paper, specifically in the context of the species *Severe acute respiratory syndrome-related coronavirus.*

To appreciate the difference between *Severe acute respiratory syndrome-related coronavirus* and SARS-CoV, i.e. between species and virus, it may be instructive to look at their relation in the context of the full taxonomy structure of several coronaviruses and in comparison with the taxonomy of the virus host, specifically humans (**Fig. 1**). Thus, SARS-CoV-Urbani with a particular genome sequence^20^ could be regarded as equivalent to a single human being, while the species *Severe acute respiratory syndrome-related coronavirus* would be on a par with the species *Homo sapiens*. This parallel could go beyond semantics and be biologically meaningful because of how coronaviruses are assigned to species in practice, although the extension of this concept to virology is yet to be developed and thoroughly tested^21^.

**Fig. 1.**
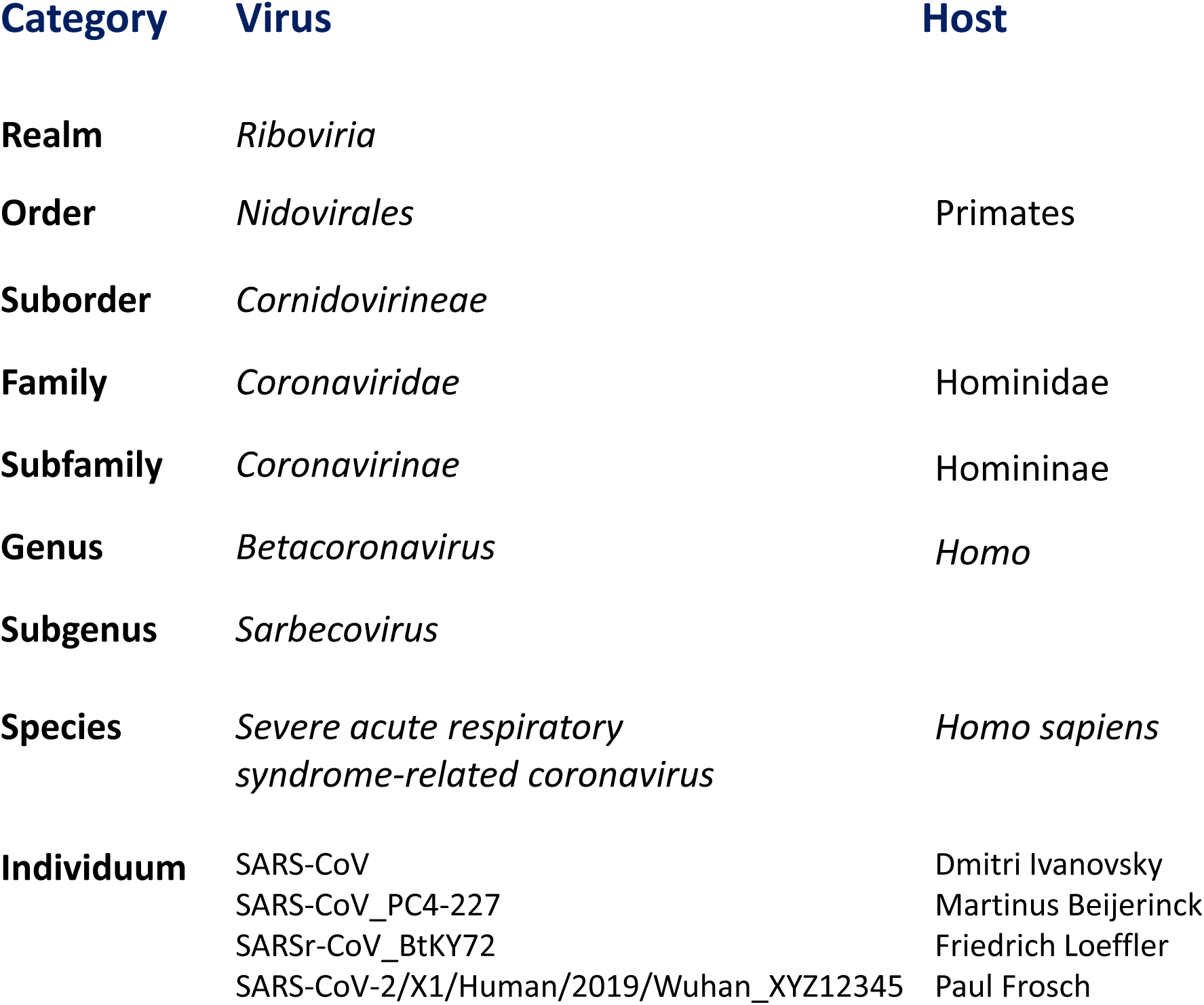
Taxonomies of coronaviruses and humans. Shown is a comparison of the taxonomies of selected coronaviruses and the founders of virology for the shared taxonomic categories. Note that these two taxonomies are independently developed using completely different criteria.

Even without knowing anything on the species concept of classifying different forms of life, every human recognizes another human as being a member of the (same) species *Homo sapiens*. However, for assigning individual living organisms to most other species, specialized knowledge and tools for assessing inter-individual differences are required. The CSG uses a computational framework of comparative genomics^22^ that is shared by several Study Groups concerned with the classification and nomenclature of the order *Nidovirales* and coordinated by the Nidovirales Study Group^23^ (**Box 3**). The Study Groups quantify and partition the variation in the most conserved replicative proteins encoded in open reading frames 1a and 1b (ORF1a/1b) of the coronavirus genome (**Fig. 2A)** to identify thresholds on pair-wise patristic distances (PPD) that demarcate virus clusters at different ranks.

**Fig. 2.**
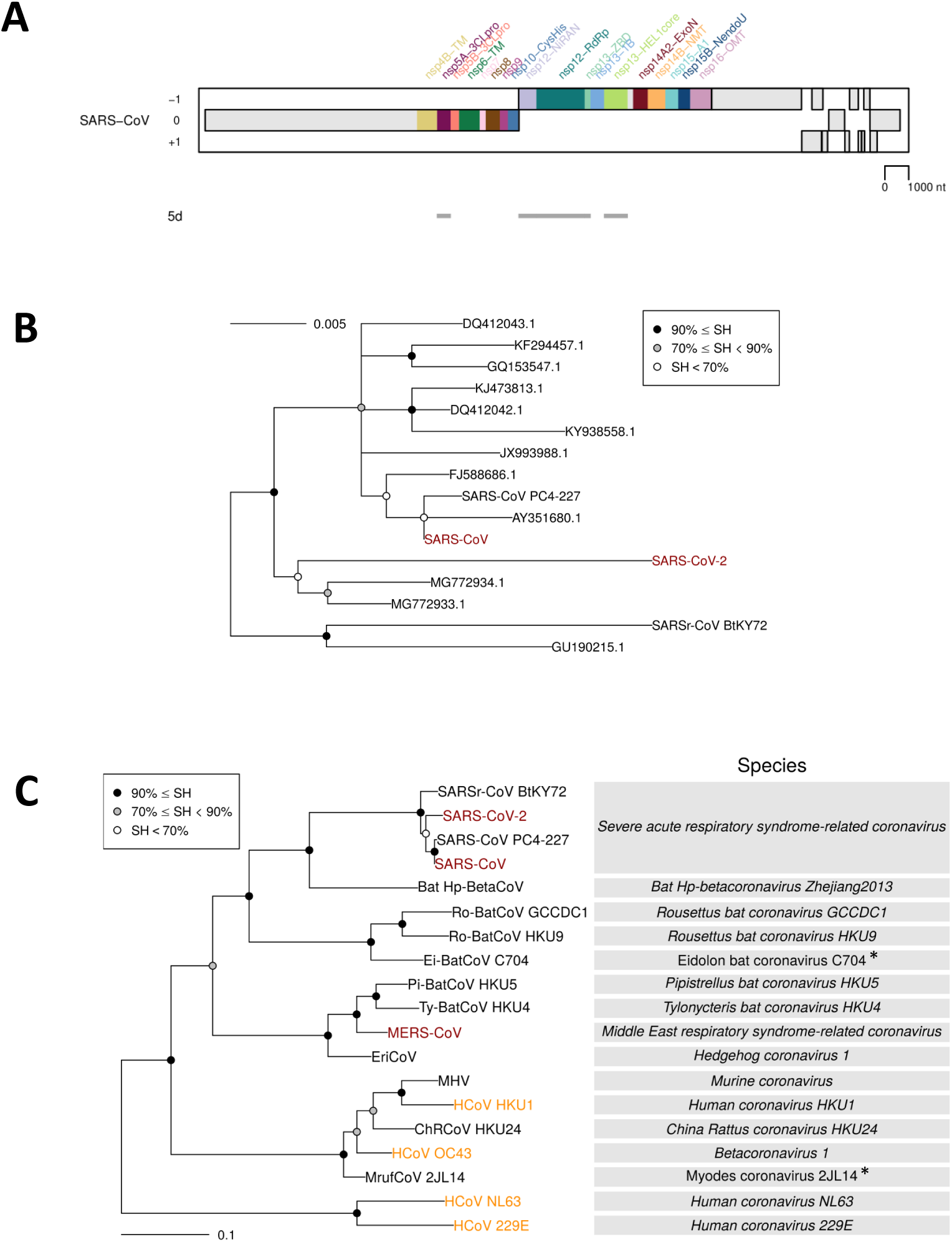
Phylogeny of coronaviruses. (**A**) Concatenated MSAs of the protein domain combination used for phylogenetic and DEmARC analyses of the *Coronaviridae* family. Shown are the locations of the replicative domains conserved in the *Nidovirales* order (5d, 5 domains: 3CLpro, 3C-like protease; NiRAN, nidovirus RdRp-associated nucleotidyltransferase; RdRp, RNA-dependent RNA polymerase; HEL1, superfamily 1 helicase with upstream Zn-binding domain (ZBD)) in relation to several other ORF1a/b-encoded domains and other major open reading frames in the SARS-CoV genome. (**B**) The Maximum-Likelihood (ML) tree of the species *Severe acute respiratory syndrome-related coronavirus* was reconstructed by IQ-Tree 1.6.1 using 83 sequences with the best fitting evolutionary model. Subsequently, the tree was purged from the most similar sequences and midpoint-rooted. Branch support was estimated using the Shimodaira-Hasegawa-like approximate likelihood ratio test (SH-aLRT) with 1000 replicates. GenBank IDs for all viruses except four are shown; SARS-CoV, AY274119.3; SARS-CoV-2 MN908947.3; SARSr-CoV_BtKY72, KY352407.1; SARS-CoV_PC4-227, AY613950.1. (**C**) Shown is an IQ-Tree ML tree of single virus representatives of thirteen species and four representatives of the species *Severe acute respiratory syndrome-related coronavirus*, genus *Betacoronavirus*. The tree is rooted with HCoV-NL63 and HCoV-229E, representing two species of the genus *Alphacoronavirus*. Red, zoonotic viruses with varying pathogenicity in humans; orange, common respiratory viruses that circulate in humans. Asterisk, ICTV approval for the two coronavirus species with non-italicized names is pending.

SARS-CoV-2 clusters with SARS-CoVs in trees of the species *Severe acute respiratory syndrome-related coronavirus* (**Fig. 2B)** and genus *Betacoronavirus* (**Fig. 2C)**, as was also reported by others^24-26^. Distance estimates between SARS-CoV-2 and the most closely related coronaviruses vary among different studies, depending on the choice of measure (nucleotide or amino acid) and genome region. Accordingly, researchers are split about the exact taxonomic position of 2019-nCoV (i.e., SARS-CoV-2). When we included SARS-CoV-2 in the dataset, including 2505 coronaviruses and used for the most recent update (May 2019) of the coronavirus taxonomy that is currently being considered by ICTV^18^, the species composition was not affected and the virus was assigned to the species *Severe acute respiratory syndrome-related coronavirus*, as detailed below.

The species demarcation threshold/limit in the family *Coronaviridae* is defined/imposed by viruses whose PPD may cross the inter-species demarcation threshold. Due to their minute share of ∼10^−4^ of the total number of all intra- and inter-species PPDs, they may not even be visually recognized in a conventional diagonal plot clustering viruses on species basis (**Fig. 3A**). Furthermore, these violators do not involve any virus of the species *Severe acute respiratory syndrome-related coronavirus* species, as evident from the analysis of maximal intraspecies PPDs of 2505 viruses of all 49 coronavirus species (**Fig. 3B)** and PDs of 256 viruses of this species **(Fig. 4**). Thus, the genomic variation of the known viruses of this species is smaller compared to that of other comparably well sampled species, e.g. those prototyped by MERS-CoV, HCoV-OC43 and IBV (**Fig. 3B**), and this species is well separated from other known coronavirus species in the sequence space. Both these characteristics of the species *Severe acute respiratory syndrome-related coronavirus* facilitate the unambiguous species assignment of SARS-CoV-2 to this species.

**Fig. 3.**
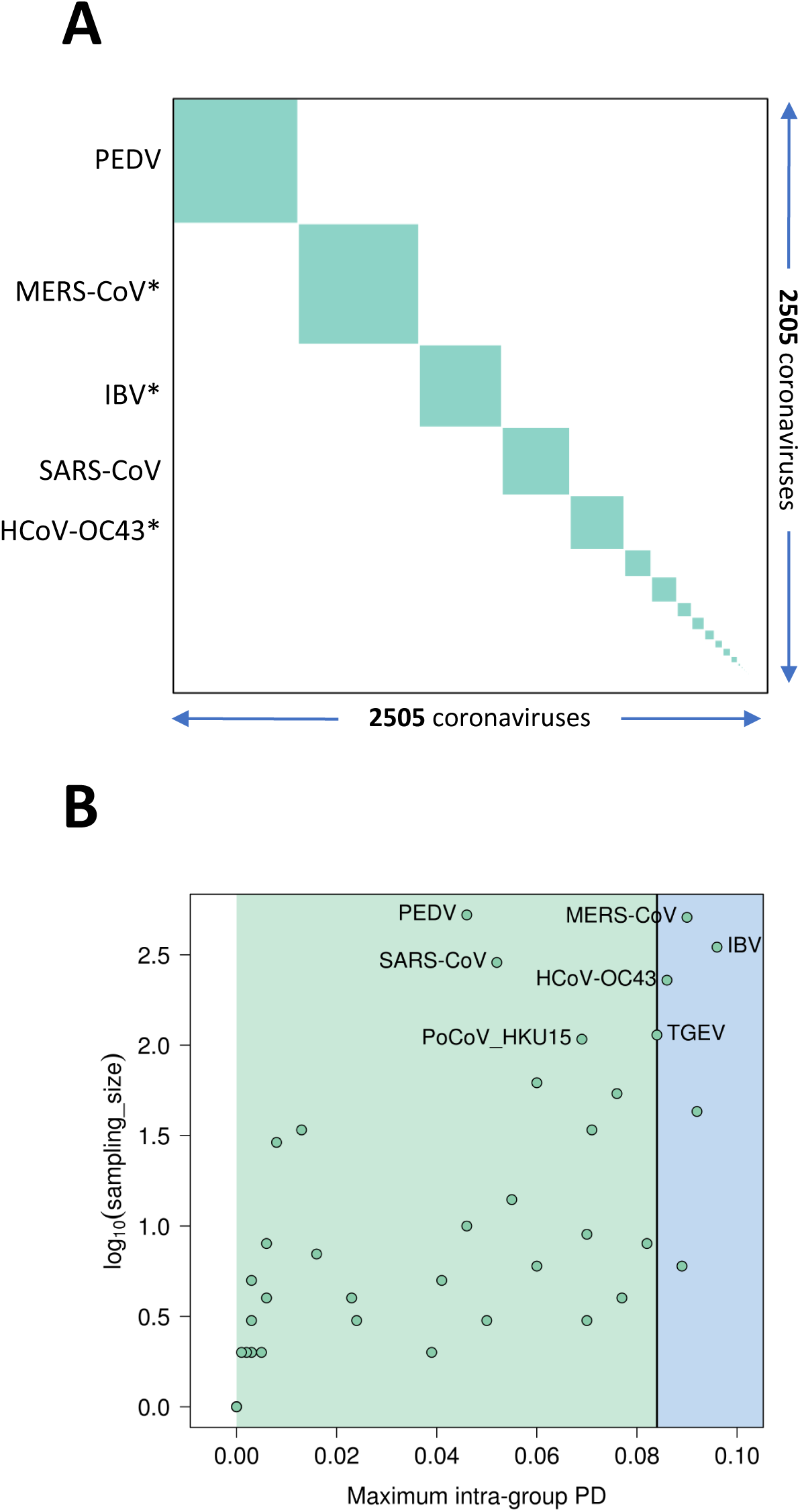
Species pairwise distance demarcation in the family *Coronaviridae*. (**A**) Diagonal matrix of PPDs of 2505 viruses clustered according to 49 coronavirus species and ordered from the most to the least populous species, from left to right; green and white, PPDs smaller and larger than the inter-species threshold (panel B), respectively. Areas of the green squares along the diagonal are proportional to the virus sampling of the respective species, and virus prototypes of the five most sampled species are specified to the left; asterisk, selected species including viruses whose some PPDs crossed threshold (“violators”). Violators of the inter-species threshold appear as white dots on the green squares along the diagonal and green dots off the diagonal, respectively; as there are just 656 dots of this kind (out of a total of 6,275,025 dots) in the panel, they may not even be visible; this is an indication of the strong support for intra-species virus clustering. (**B**) Maximal intra-species PPDs (X axis, linear scale) plotted against virus sampling (Y axis, log scale) for 49 species (green dots) of the *Coronaviridae*. Indicated are the acronyms of virus prototypes of the seven most sampled species. Green and blue plot sections, intra-species and intra-subgenera PPD ranges. Vertical black line, inter-species threshold.

**Fig. 4.**
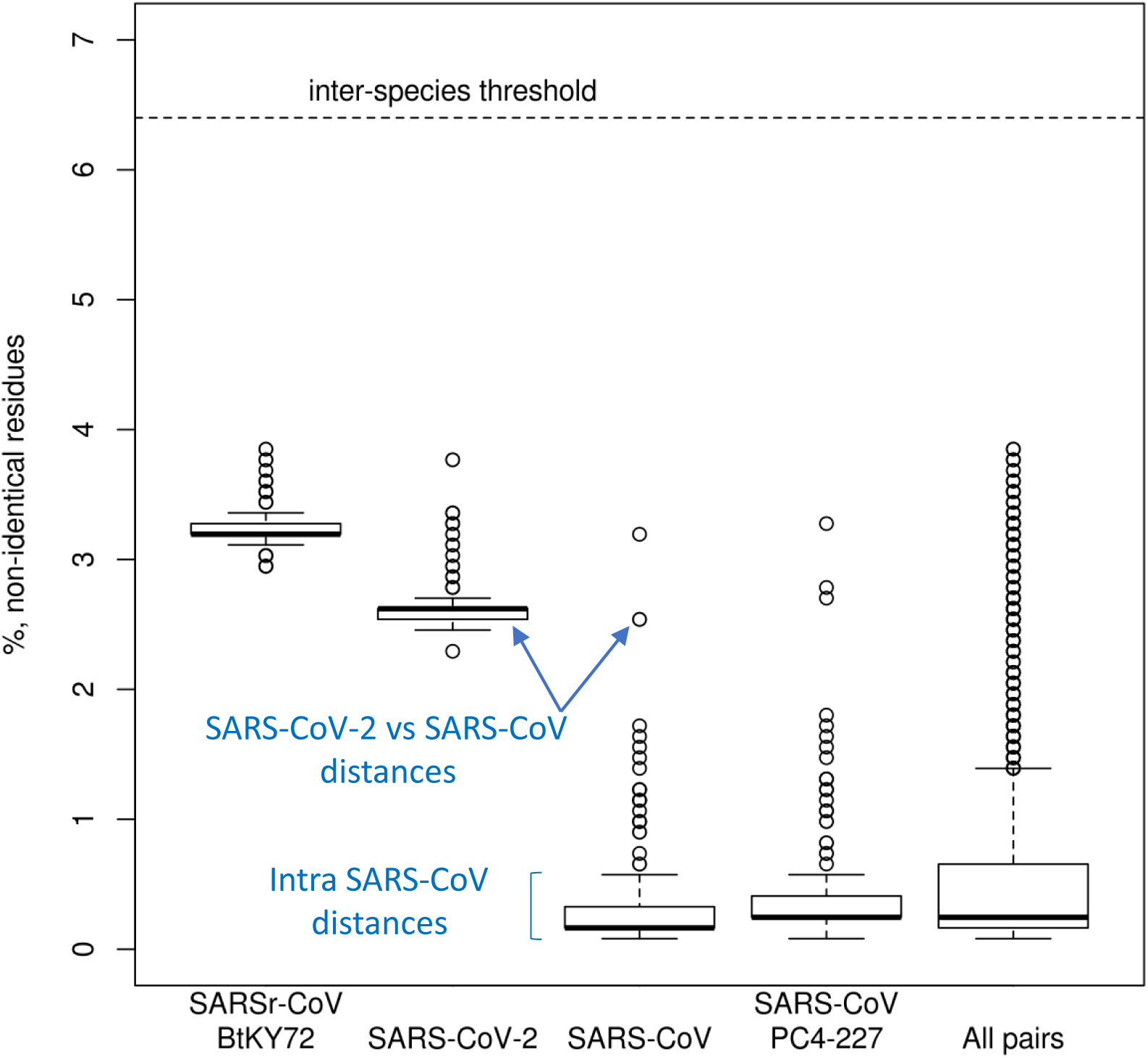
Pairwise distances between selected viruses within the species *Severe acute respiratory syndrome-related coronavirus.* Shown are the PDs in % of non-identical residues (Y axis) for four viruses representing three major phylogenetic lineages of the species (Fig. 2B) and all pairs of the 256 viruses of this species (“All pairs”). The PD values were derived from the PPD values (Fig. 3).

Intra-species PDs of SARS-CoV-2 belong to the top 25% of this species and also include the largest PD, that between SARS-CoV-2 and an African bat virus isolate (SARSr-CoV_BtKY72)^27^ (**Fig. 4**), representing two basal lineages within the species *Severe acute respiratory syndrome-related coronavirus* that constitute very few known viruses (**Fig. 2BC**). These relationships stand in contrast to the shallow branching of the most populous lineage of this species which includes all the human SARS-CoV isolates collected during the 2002-2003 outbreak and the closely related bat viruses of Asian origin identified in the search for the potential zoonotic source of that epidemic^28^. (Note that this clade structure is susceptible to homologous recombination, which is common in this species^29 28,30^; to formalize clade definition, it must be revisited after the virus sampling of the deep branches was improved sufficiently). The current sampling defines a very small median PD for human SARS-CoVs, which is approximately 15 times smaller than the median PD determined for SARS-CoV-2 (0.16% vs 2.6%, **Fig. 4**). This small median PD of human SARS-CoVs also dominates the species-wide PD distribution (0.25%, **Fig. 4**). Along with the initial failure to detect the causative agent of the disease using SARS-CoV-specific PCR setups, the separation from SARS-CoV in the phylogeny and the PD space explains why 2019-nCoV (SARS-CoV-2) may be considered a novel virus by many researchers.

## Designating 2019-nCoV as SARS-CoV-2 and providing guidance for naming its variants

The above results show that, in terms of taxonomy, SARS-CoV-2 is (just) another virus in the species *Severe acute respiratory syndrome-related coronavirus*. In this respect, the discovery of this virus differs considerably from the description of the two other zoonotic coronaviruses, SARS-CoV and MERS-CoV, introduced to humans in the 21^st^ century (**Fig. 5A**). Both these viruses were considered novel by this study group based on prototyping two species and two informal subgroups of the *Betacoronavirus* genus that were recently recognized as subgenera *Sarbecovirus* and *Merbecovirus*^17,31,32^. Due to being first, these viruses and their taxa were assigned new names whose origins reflected the practice and the state of virus taxonomy at the respective times (**Fig. 5B**) (**Box 4**). Neither of these circumstances are applicable to SARS-CoV-2, which is assigned to an existing species of hundreds of known viruses predominantly isolated from humans and diverse bats. All these viruses have names derived from SARS-CoV (directly or through the species name), even though only the human isolates collected during the 2002-2003 outbreak have been confirmed to cause SARS in infected individuals. Thus, the reference to SARS in all these virus names (combined with the use of specific prefixes, suffixes and/or genome sequence IDs in public databases) acknowledges the phylogenetic grouping of the respective virus with viruses isolated from SARS patients, for example SARS-CoV-Urbani, rather than linking this virus to a specific disease (i.e., SARS) in humans. Based on the established practice of virus naming in this species and the relatively distant relationship of SARS-CoV-2 to the prototype SARS-CoV in a species tree and the distance space (**Figs. 2B**, and **4**), the CSG renames 2019-nCoV to SARS-CoV-2.

**Fig. 5.**
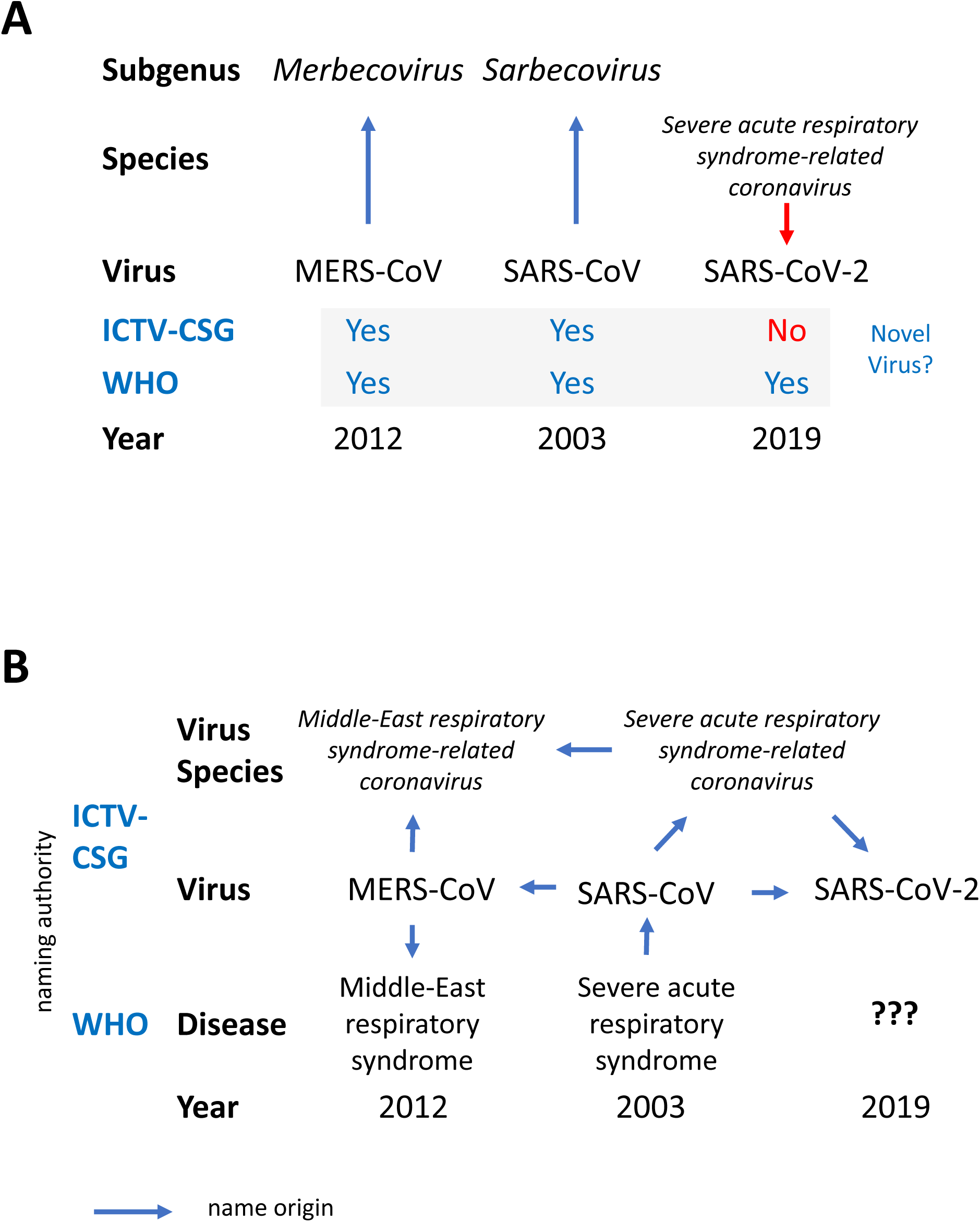
Virus novelty and naming of the three zoonotic coronaviruses emerging in the first decades of the 21^st^ century. Year, indicates the year in which the virus was first identified. (**A)** Independent assessments of virus novelty by the ICTV-CSG and WHO performed during the three outbreaks came to different conclusions. Vertical arrows indicate the degree of virus novelty according to taxonomy. (**B**) History of coronavirus naming during the three zoonotic outbreaks in relation to virus taxonomy and disease (clinical manifestation).

In contrast to SARS-CoV, the name SARS-CoV-2 has NOT been derived from the name of the SARS disease (**Fig. 5B**), and in no way, it should be used to predefine the name of the disease (or spectrum of diseases) caused by SARS-CoV-2 in humans, which will be decided upon by the WHO. The available yet limited epidemiological and clinical data for SARS-CoV-2 suggest that the disease spectrum, and transmission modes of this virus and SARS-CoV may differ^1^. Also, the diagnostic methods used to confirm SARS-CoV-2 infections are not identical to those of SARS-CoV. This is reflected by the specific recommendations for public health practitioners, healthcare workers and laboratory diagnostic staff for SARS-CoV-2/2019-nCoV (e.g. WHO guidelines for 2019-nCoV; https://www.who.int/emergencies/diseases/novel-coronavirus-2019). By uncoupling the naming conventions used for coronaviruses and the diseases they may cause in humans and animals, we wish to help the WHO with naming diseases in the most appropriate way (WHO guidelines for disease naming; https://apps.who.int/iris/handle/10665/163636) (**Fig. 5B**).

To facilitate good practice and scientific exchange, the CSG recommends that researchers describing new isolates of this virus and other viruses in this species adopt a standardized format for public databases and publications. The proposed naming convention includes a reference to the host organism that the virus was isolated from, the time of isolation, and the place of isolation (geographic location): Virus/Isolate/Host/Date/Location, e.g. SARS-CoV-2/X1/Human/2019/Wuhan. This complete designation along with additional and important characteristics, such as association with pathogenicity in humans or other hosts, should be included in the submission of each isolate genome sequence to public databases, e.g. GenBank. In publications, this name could be further extended with a sequence database ID, e.g. SARS-CoV-2/X1/Human/2019/Wuhan_XYZ12345, when first mentioned in the text. We believe that this format will inform about the major characteristics of each particular virus isolate (genome sequence) that are critical for subsequent epidemiological and other studies, as well as control measures.

## Concluding remarks: from focusing on pathogens to understanding virus species

Historically, public health and fundamental research have been focused on the detection, containment, treatment and analysis of viruses that are pathogenic to humans, with little regard to exploring and defining their genetic diversity and biological characteristics as a species. In this framework, the emergence of SARS-CoV-2 as a human pathogen in December 2019 may be perceived as completely independent from the SARS-CoV outbreak in 2002-2003. Although SARS-CoV-2 is NOT a descendent of SARS-CoV **(Fig. 2B)** and the introduction of each of these viruses into humans was likely facilitated by unknown external factors, the two viruses are genetically so close to each other **(Fig. 2C)** that their evolutionary histories and characteristics are mutually informative. Our understanding of these pathogens could be significantly advanced if both viruses were characterized along with viruses of other origins, known and yet-to-be discovered^25^, as part of the *Severe acute respiratory syndrome-related coronavirus* species, with the long-term goal of comprehending the biology and evolution of that species, as is the norm elsewhere in biology. To connect this development to health care, diagnostic tools that target the entire species should complement existing tools that detect individual pathogenic variants.

Although this paper focuses on a single virus species, the raised issues concern other species in the family and possibly beyond. As a first step toward appreciation of this species and its cousins, researchers, journals, databases, and other relevant bodies should adopt proper referencing to the full taxonomy of coronaviruses under study. This includes that the relevant virus species is explicitly acknowledged along with the viruses included in this species by following the ICTV naming rules (**Box 4**) which, regretfully, are rarely observed in common practice, contributing to the proliferation of mixing viruses and species in the literature (and the authors of this paper wish to acknowledge that they were also not immune to this problem in several cases). This necessary adjustment may be facilitated by the major revision of the virus species nomenclature that is currently being discussed by ICTV and planned to be implemented in the near future^33^. With this change in place, the CSG is resolved to address the existing significant overlap between virus and species names that complicates the appreciation and use of the species concept in its application to coronaviruses.

**Box 1**

#### Virus Discovery: from disease-based to phenotype-free

Understanding the cause of a specific disease that spreads among individuals of the same biological species (infectivity) was the major driving force for the discovery of the first virus, initially in plants, and many others in all types of life, including humans. The range of diseases and hosts that specific viruses were confirmed to be associated with have been the two key and most appreciated characteristics used to define viruses that are invisible to the naked eye due to their minute size^34^. They belong to the so-called phenotype of viruses, which includes those that – like a disease – are shaped by virus-host interactions, e.g. transmission rate or immune correlates of protection, and others that are virus-specific, e.g. the architecture of virus particles. These phenotypic features are of critical importance for many decisions and actions related to medically and economically important viruses, especially during outbreaks of severe infectious diseases, and they dominate the general perception of viruses.

However, the host is not definitive nor is the pathogenicity known for a major (and fast growing) share of viruses, including many coronaviruses discovered in metagenomics studies using next generation sequencing technology^35,36^. These studies analyze diverse environmental specimens and assemble genomic sequence of viruses, which circulate in nature and have never been characterized on the phenotypic level. Thus, the genome sequence is the only characteristic that is known for the vast majority of viruses, and its use in defining virus identity in the virosphere is the only available choice going forward. In this framework, a virus is defined by its genome sequence that instructs the synthesis of polynucleotide molecules capable of autonomous replication inside cells and dissemination between cells or organisms under appropriate conditions. It may or may not be harmful to its natural host. Experimental studies may be performed for a fraction of known viruses, while computational comparative genomics is used to classify (and deduce characteristics of) all viruses.

**Box 2**

#### Recognizing virus novelty

Besides haplotypes of a virus quasispecies, the terms strains and isolates are in common use to refer to virus variants with larger genome variations, although there are different opinions as to which term should be used in a specific context. If a candidate virus clusters within a group of isolates, it is a variant of this group and, in other words, may be considered a known virus. On the other hand, if the candidate virus is outside of known groups and its distances to viruses of these groups are comparable to those observed between viruses of different groups (intergroup distances), the candidate virus is distinct and could be considered novel. Commonly this evaluation is conducted *in silico* using phylogenetic analysis that may be complicated by uneven rates of evolution that vary across different virus lineages and genomic sites due to mutation, including exchange of genome regions in closely related viruses (homologous recombination). Comparative genomics forms also the basis for PCR assays that are suitable to detect established viruses and their groups *in vitro*. If such PCRs do not recognize the candidate virus, it may be considered novel. There are two caveats to the above approaches. Since the current sampling of viruses is small and highly biased toward viruses of significant medical and economic interest, the group composition varies tremendously among different viruses, making decisions on novelty group-specific and dependent on the choice of specific criteria selected by researchers. Practically, this means that definitive evidence for novelty in one group may not stand up to scrutiny in another.

The challenges mentioned above are addressed specifically in the framework of virus taxonomy, which partitions genomic variation above strain/isolate level and develops a unique taxa nomenclature under the supervision of the ICTV^21,37^. To decide on virus novelty, taxonomists use the results of different analyses, although comparative sequence analysis plays an increasing role and is now the primary tool in the classification of coronaviruses (**Box 3**). Taxonomical classification is hierarchical, using nested groups (taxa) that populate different levels (ranks) of classification. Taxa of different ranks differ in respect to intra-taxon pairwise divergence, which increases from the smallest at the species rank to the largest at the realm rank. They may also be distinguished by taxon-specific markers that characterize natural groupings. When classifying a virus, researchers are required to define and name taxa only on the species and genus ranks while filling other ranks is optional. Thus, species is the smallest and mandatory unit of virus taxonomy. Only if a virus prototypes a new species, it will be regarded as truly novel, taxonomy-wise. Within this framework, a virus that crosses a host barrier and acquires novel properties remains part of the original species. This association may persist even after the virus established a permanent circulation in the new host, as it likely happened with coronaviruses of four species circulating in humans (reviewed in^38^).

**Box 3**

#### Classifying coronaviruses

In the past, the classification of coronaviruses was largely based on serologic (cross-) reactivity involving the S protein till it became based on comparative sequence analysis of replicative proteins. The choice of proteins and the methods used to analyze them have gradually evolved since the start of this century^19,31,32,39^. Currently, the CSG analyzes 3CLpro, NiRAN, RdRp, ZBD and HEL1^40^ (**Fig. 2A**), which replaced the seven domains used for analysis between 2009 and 2015^17^. According to our current knowledge, these five most essential domains are the only domains that are conserved in all viruses of the order *Nidovirales;* they are used for the classification by all nidovirus SGs (coordinated by the NSG).

Since 2011, the classification of coronaviruses and other nidoviruses has been assisted by the DEmARC (DivErsity pArtitioning by hieRarchical Clustering) software which defines taxa and ranks^22^. Importantly, the involvement of all coronavirus genome sequences available at the time of analysis allows family-wide designations of demarcation criteria for all ranks, including species, regardless of the taxa sampling size, be it a single or hundreds of virus(es). DEmARC delineates monophyletic clusters (taxa) of viruses, using weighted linkage clustering in the pairwise patristic distance (PPD) space and according to the classification ranks defined through clustering cost (CC) minima presented as PPD thresholds. The persistence of thresholds in the face of increasing virus sampling is interpreted in the DEmARC framework as a reflection of biological forces and environmental factors^41^. Specifically, homologous recombination, which is common in coronaviruses^42-44^, is believed to be restricted in the most essential proteins, like those used for classification, to within a species. This restriction promotes intra-species diversity and contributes to inter-species separation; hence, they are biological entities, which deviates from the current ICTV definition of virus species as man-made constructs ^21^. To facilitate the use of rank thresholds outside the DEmARC framework, they are converted into pair-wise differences (PD) %, which researchers commonly use to arrive at a tentative assignment of a given virus within the coronavirus taxonomy following conventional phylogenetic analysis of selected viruses.

**Box 4**

#### Naming Viruses and Virus Species: roles of ICTV and WHO

Besides humans and their pets, viruses may be the only biological entities that have names for virtually every single known representative, in addition to the groups (taxa) that together constitute the official classification of virus taxonomy. This exceptional treatment of viruses is a by-product of the historical perception of viruses as a feature of the diseases they cause (**Box 1**) and the way we usually catalogue and classify newly discovered viruses, rather than an expression of appreciation of virus individuality. Apart from disease, also geography and the organism a given virus was isolated from dominate the name vocabulary, occasionally engraving connections that may be accidental (rather than typical) to the virus in nature. Virus naming is linked to the recognition of virus novelty (**Box 2**), which is not formally regulated; and no national or global authorities have been established to certify virus novelty and approve virus names. Thus, it is mainly up to virus discoverers or other researchers to decide on these matters, and it is not uncommon to see two variants of the same virus having very different names, if they were described by different researchers.

One of the priorities of the World Health Organization (WHO), an agency of the United Nations, is concerned with communicable diseases, including the coordination of international public health activities aimed at containing (and mitigating the consequences of) major virus epidemics. WHO also considers virus novelty and has responsibility in naming the disease(s) caused by newly emerging human viruses. By doing so, WHO often takes the traditional approach of linking viruses to specific diseases (**Box 1**) and assessing novelty by an apparent failure to detect the causative agent by established diagnostic assays. Formally, ICTV was involved in virus naming only until the grouping of the most closely related viruses into clusters (taxa species) was introduced and their nomenclature in the taxonomy established ^45^. Since then, the specialized Study Groups have been involved in virus naming only on a case-by-case basis as an extension of their official remit and using the special expertise of their members. However, since the study groups are responsible for assigning viruses to virus species, they could play an important role in advancing the species concept (which is yet to be fully appreciated) to the research community and others, who may be uncertain about its significance ^21^. The CSG adopts the DEmARC-based approach to define virus species (**Box 3**). Practically, virus species are often ignored or confused with viruses that are part of the respective species. This problem may be alleviated by strictly adhering to established rules for the naming of species and viruses in that species ^33^. The species name is italicized, starts with a capital letter, and must NOT be spelled in an abbreviated form; neither of these rules and conventions apply to virus names, hence severe acute respiratory syndrome coronavirus, or SARS-CoV as it is widely known.

## AUTHOR CONTRIBUTIONS

Susan C. Baker, Ralph S. Baric, Christian Drosten, Raoul J. de Groot (RDG), Alexander E. Gorbalenya (AEG), Bart L. Haagmans, Benjamin Neuman (BN), Stanley Perlman, Leo L. M. Poon, Isabel Sola, and John Ziebuhr (JZ), are members of the Coronavirus Study Group (CSG), chaired by JZ; RDG, AEG, Chris Lauber (CL), BN, and JZ are members of the Nidovirales Study Group (NSG), chaired by AEG; AEG and JZ are members of the International Committee on Taxonomy of Viruses. AEG, Anastasia A. Gulyaeva (AAG), CL, Andrey M Leontovich, Dmitry Penzar, Dmitry Samborskiy (DS), and Igor A. Sidorov are members of the DEmARC team led by AEG. DS generated the classification of SARS-CoV-2 using a computational pipeline developed by AAG and using software developed by the DEmARC team. The CSG considered and approved this classification. AEG and JZ drafted and revised the text. AEG and DS generated the figures. All authors reviewed the manuscript and approved its submission for publication.

## ACKNOWLEDGMENTS

The authors gratefully acknowledge the work of all researchers who released SARS-CoV-2 genome sequences through the GISAID initiative and particularly the authors of the MN908947 genome sequence. AEG is indebted to members of the ICTV Executive Committee for discussions of classification and nomenclature issues relevant to this paper.

